# Animal biosynthesis of complex polyketides in a photosynthetic partnership

**DOI:** 10.1101/764225

**Authors:** Joshua P. Torres, Zhenjian Lin, Jaclyn M. Winter, Patrick J. Krug, Eric W. Schmidt

## Abstract

Animals are rich sources of complex polyketides, including pharmaceuticals, cosmetics, and other products. Most polyketides are associated with microbial or plant metabolism^1^. For this reason, symbiotic bacteria or dietary organisms are often the true producers of compounds found in animals^2,3^. Although increasing evidence suggests that animals themselves make some compounds^4–7^, the origin of most polyketides in animals remains unknown. This problem makes it difficult to supply useful animal compounds as drugs and severely constrains our understanding of chemical diversity and the scope of biosynthesis in nature. Here, we demonstrate that animals produce microbe-like complex polyketides. We report a previously undocumented but widespread branch of fatty acid synthase- (FAS)-like proteins that have been retooled by evolution to synthesize complex products. One FAS-like protein uses only methylmalonyl-CoA as a substrate, otherwise unknown in animal lipid metabolism, and is involved in an intricate partnership between a sea slug and captured chloroplasts. The enzyme’s complex, methylated polyketide product results from a metabolic interplay between algal chloroplasts and animal host cells, and also likely facilitates the survival of both symbiotic partners, acting as a photoprotectant for plastids and an antioxidant for the slug^8–12^. Thus, we find that animals can unexpectedly synthesize a large and medically useful class of structurally complex polyketides previously ascribed solely to microbes, and can use them to promote symbiotic organelle maintenance. Because this represents an otherwise uncharacterized branch of polyketide and fatty acid metabolism, we anticipate a large diversity of animal polyketide products and enzymes awaiting discovery.

## MAIN

Complex, branched polyketides, are canonically ascribed to microbial metabolism, yet they are commonly found in animals^13,14^. The sacoglossan polypropionate pyrones are among the most structurally complex polyketides isolated from animals (Fig. 1). Sacoglossans consume algae, and some species maintain the stolen algal chloroplasts (kleptoplasts) for weeks or in exceptional cases, for several months^15^. Feeding studies showed that fixed carbon obtained from *de novo* photosynthesis is efficiently incorporated into polypropionates via chloroplasts^16^. Labeled propionate is also incorporated^17–20^, suggesting that the mollusk compounds are made via the methylmalonate pathway, which has historically been associated with bacterial metabolism^24^. The polypropionates react via oxidative and photocyclization pathways that are likely non-enzymatic to form the mature natural products^8–10,16^. It is thought that these reactions protect the slugs from damage associated with photosynthesis, and that they may thus be necessary for a photosynthetic lifestyle^8–12^. Thus, the polypropionates are known to originate in the mollusks and to be central to mollusk biology, yet their biochemical sources were unknown.

**Fig. 1:**
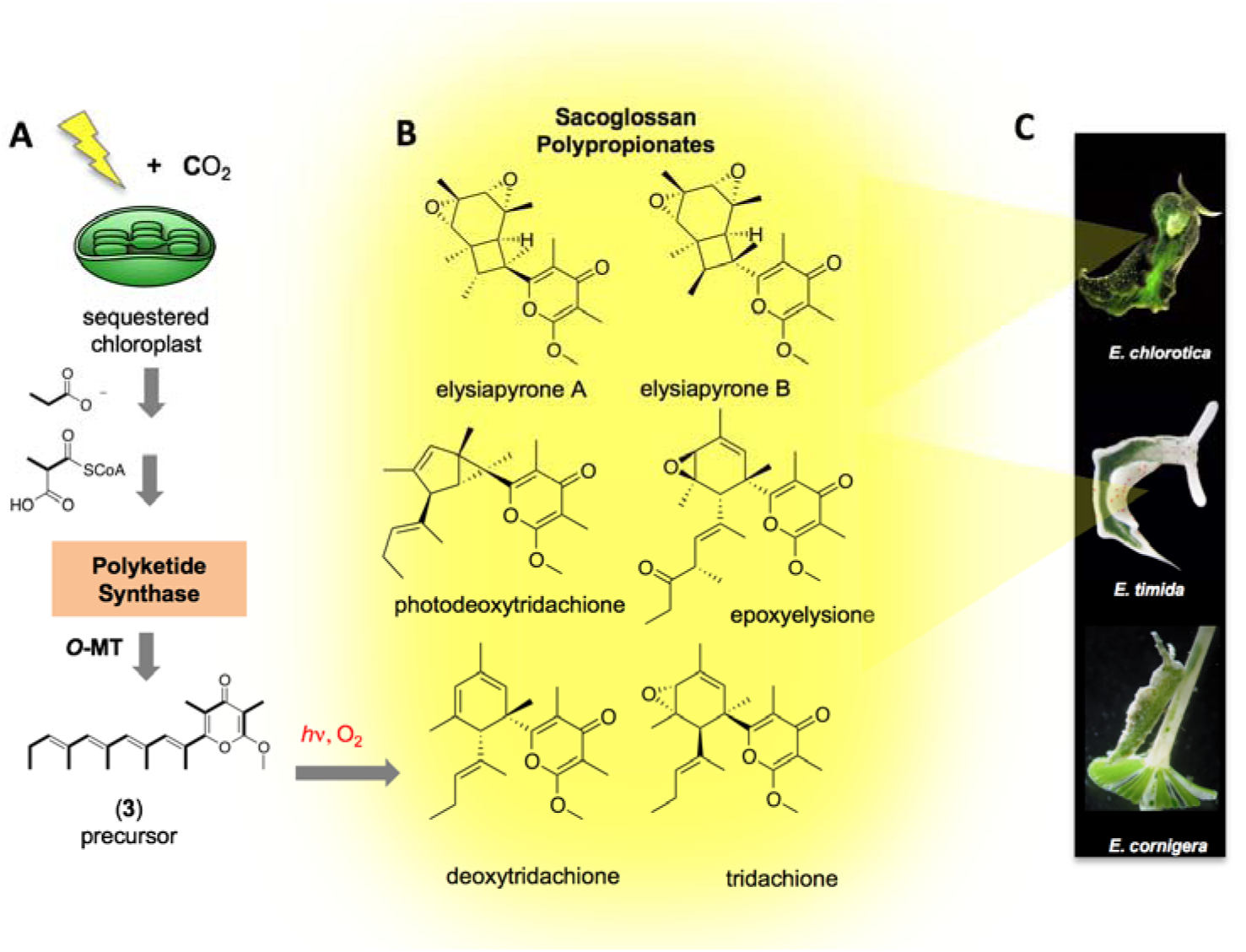
Biogenesis of sacoglossan polypropionates. **a** Previous feeding studies demonstrated that kleptoplasts in the sacoglossans fix carbon that is used to make polypropionates. Propionate is also a precursor, presumably via methylmalonate. A further series of hypothetical steps, including synthesis by a PKS, methylation, and photochemical and oxidation reactions would lead to **b** the known natural polypropionates of diverse structures. These are found in **c** sacoglossan species investigated in this work, including *E. chlorotica* and *E. timida*, but not *E. cornigera*.

We took an unbiased approach to define the molecular source of polypropionates. To test the bacterial origin hypothesis, we sequenced the metagenomes of polypropionate-containing sacoglossans *Plakobranchus* cf. *ocellatus* “aff. sp. 1”^21^ and *Elysia diomedea*. We were unable to identify good candidate polypropionate synthases from the bacterial metagenomes, suggesting a possible origin in the animals or chloroplasts. We took advantage of recently released genomes and transcriptomes of three sacoglossans^11,22–24^, *Elysia chlorotica, E. timida*, and *E. cornigera*, to test this hypothesis. *E. chlorotica* contains polypropionates *ent*-9,10-deoxytridachione and elysione^25^, while *E. timida* contains *ent*-9,10-deoxytridachione and several relatives (Fig. 1, Extended Data Table 1)^26^. Both species obtain fixed carbon from chloroplasts that they maintain for one (*E. timida*) or several (*E. chlorotica*) months. In contrast, no polypropionates have been reported from *E. cornigera*, which only retains chloroplasts for a few days^27^.

Based upon recent reports of animals that make polyene fatty acids^5,6^, we suspected that the animals might encode modified fatty acid synthase (FAS) or PKS enzymes. Examining the available *E. chlorotica* transcriptomes, we found a number of fragmented transcripts encoding multiple type I FAS enzymes. Reassembly of those transcripts from raw read data revealed four different FAS-like enzymes (Extended Data Table 2). Two of these were the cytoplasmic and mitochondrial FAS enzymes of primary metabolism. The remaining two (which we named EcPKS1 and EcPKS2) formed their own distinct group in a global PKS/FAS tree that included bacterial, fungal, and animal PKS variants (Fig. 2A, Extended Data Fig. 1). *EcPKS1, EcPKS2*, and cytoplasmic *FAS* were encoded in the animal genomes, and not in chloroplasts (Fig. 2C, Extended Data Fig. 2). *EcPKS1* homologs were only found in *E. chlorotica* and *E. timida*, and not in *E. cornigera*, while the other genes were found in all three. This observation was consistent with EcPKS1 being the putative polypropionate synthase.

**Fig. 2:**
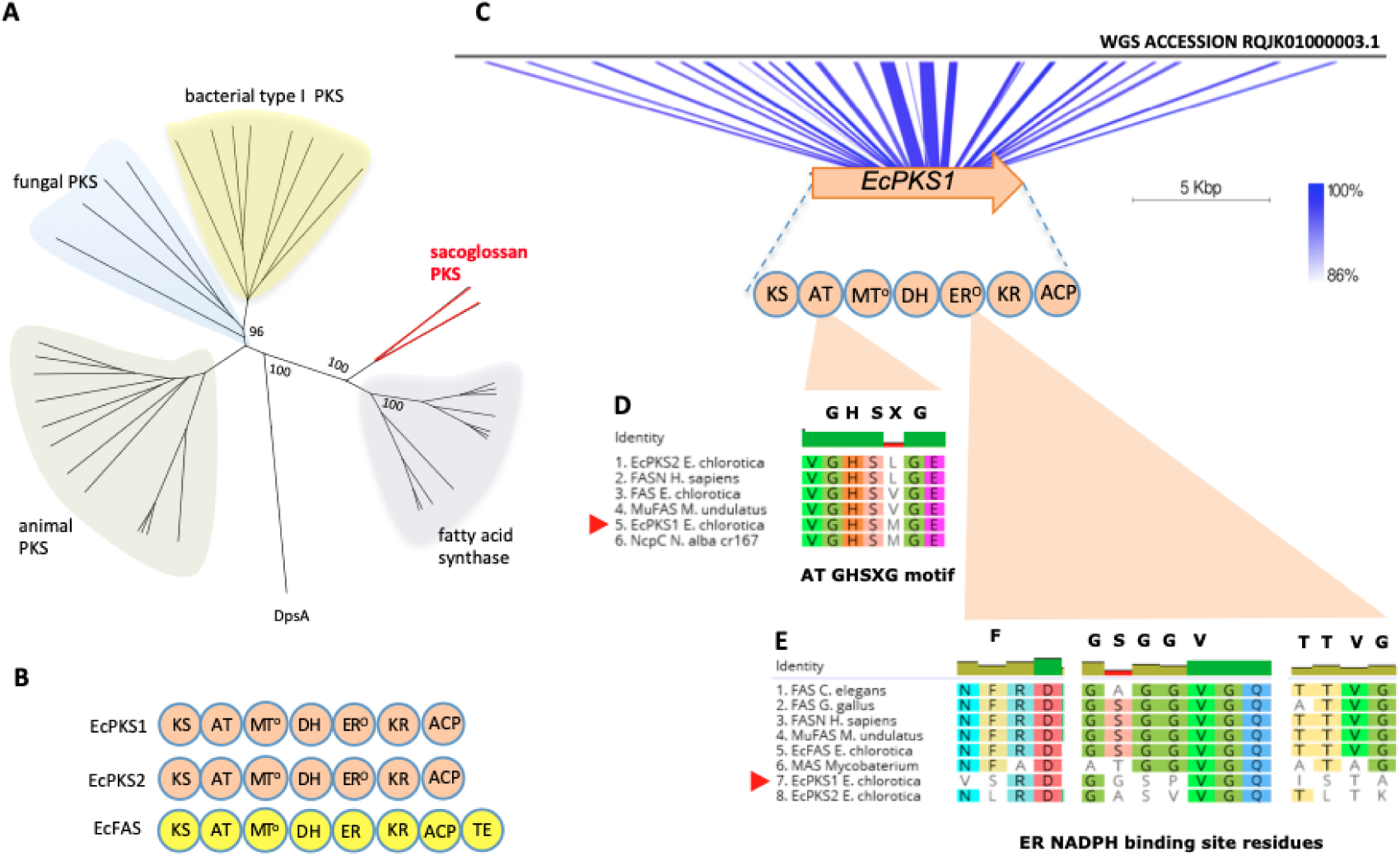
Sacoglossan PKSs, a new class of FAS-like PKS enyzmes. **a** Sacoglossan PKSs represent a group separate from the known FAS and PKS proteins from bacteria, fungi and animals. Numbers indicate consensus support in percent. Outgroup DpsA is a type II PKS sequence from bacteria, while the remainder are type I enzymes. **b** The domain architecture of the sacoglossan PKSs, EcPKS1 and EcPKS2 includes ketosynthase (KS), acyltransferase (AT), methyltransferase (MT), dehydratase (DH), enoylreductase (ER), ketoreductase (KR) and acyl carrier protein (ACP). This domain architecture is identical to the sacoglossan FAS, EcFAS except for the absence of a C-terminal thioesterase domain. **c** *EcPKS1* is encoded in the genome of *E. chlorotica*. **d** The acyltransferase (AT) domain sequence indicates loading preference for methymalonate while the **e** enoylreductase (ER) domain lacks key NADPH binding site residues making this function inactive.

Like other type I FAS and PKS enzymes, EcPKS1 and EcPKS2 contain multiple domains responsible for product formation (Fig. 2B). Enoylreductase (ER) domains reduce double bonds in the nascent polyketide chain. Because EcPKS1 and EcPKS2 lack functional ER domains, they likely both synthesize polyenes (Fig. 2D)^5,28^. The acyltransferase (AT) domain is responsible for substrate selection, and therefore would be responsible for choosing malonyl-CoA (straight-chain lipids) or methylmalonyl-CoA (methyl-branched lipids)^29^. In bacteria, amino acid residues responsible for AT selectivity are well understood^30^. Since no sequenced animal FASs/PKSs prefer methylmalonyl-CoA, there is no model for AT selectivity in animals. Gratifyingly, EcPKS1, EcPKS2, and several relatives, contained a conserved GHSXGE sequence motif found in bacterial ATs that is known to be important in substrate selection. In bacteria, when X is methionine, the substrate is methylmalonyl-CoA. In EcPKS1, a GHSMGE sequence was observed, whereas that of EcPKS2 was GHSLGE (Fig. 2E). This implied that EcPKS1 might use methylmalonyl-CoA, while EcPKS2 should use malonyl-CoA. Thus, EcPKS1 had all of the sequence and expression features consistent with the polypropionate synthase.

*EcPKS1* gene was synthesized and expressed solubly in *Saccharomyces cerevisiae* BJ5464 harboring the *npgA* phosphopantetheinyl transferase gene (Extended Data Fig. 3)^31^. Incubation of the purified enzyme with methylmalonyl-CoA and NADPH led to a series of new peaks in the chromatographic trace (Fig. 3A, Extended Data Fig. 4). The major peak had a UV spectrum and mass consistent with a triene pyrone that is three carbons smaller than the major natural products of *E. chlorotica* (Fig. 3B). A second, minor tetraene pyrone product had mass and UV features consistent with the putative precursor of tridiachiones and related natural products^17^.

**Fig. 3.**
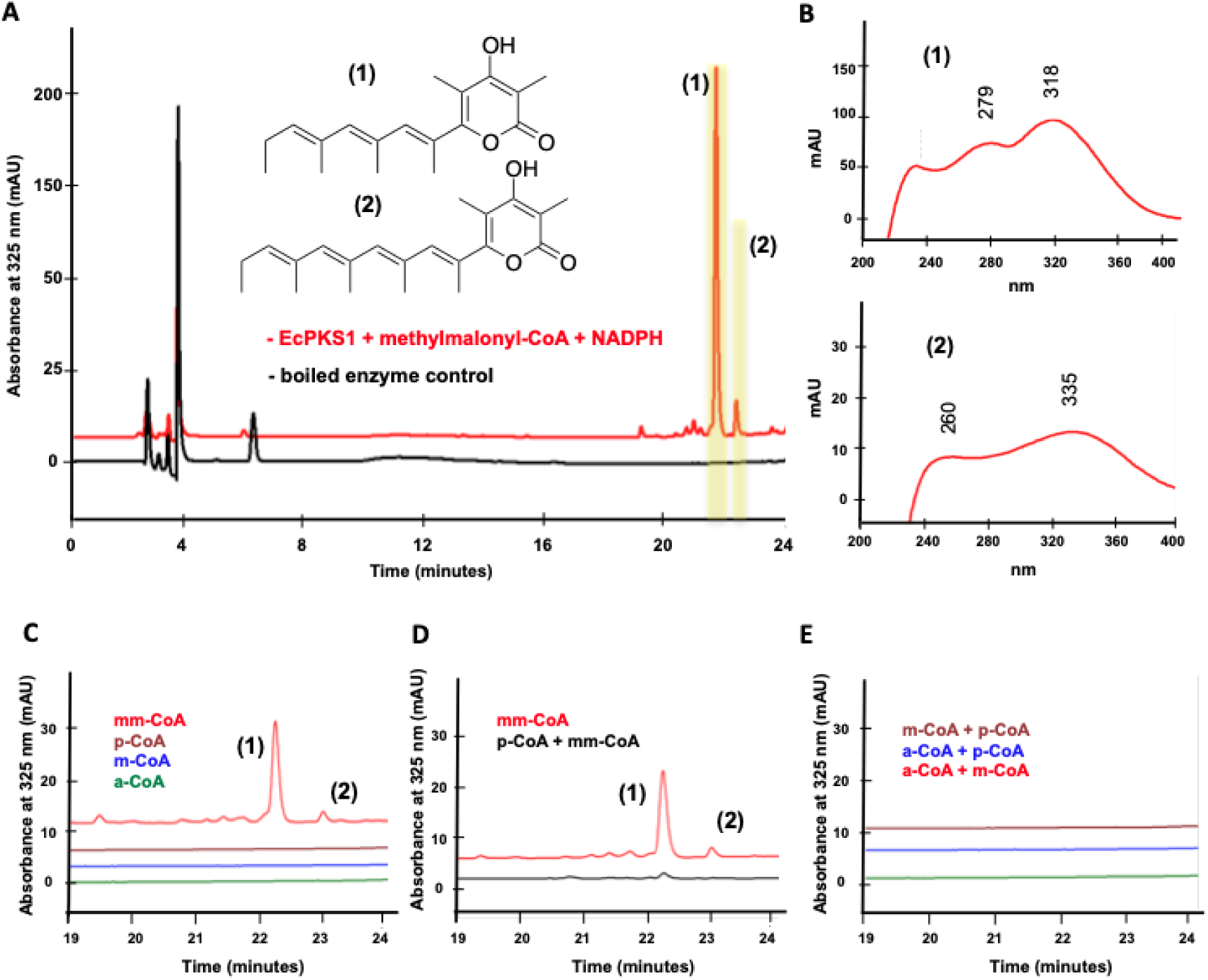
EcPKS1 is a methylmalonate-specific animal FAS/PKS that synthesizes the tridachione precursor. **a** Incubation of EcPKS1 with methylmalonyl-CoA and NADPH resulted in the synthesis of polyene **1** and tridachione precursor **2. b** UV spectra of the synthesized products showing features of polyene propionates consistent with the structural assignment. **c** Incubation with other acyl-CoA substrates (acetyl-CoA: a-CoA, malonyl-CoA: m-CoA, propionyl-CoA: p-CoA, methylmalonyl-CoA: mM-CoA) with NADPH did not yield new products. **d** Using propionyl-CoA as the loading molecule inhibited the reaction, as observed in decreased yields. **e** Using different loading and extender units other than methylmalonate did not produce any products. Graphs shown in **a** and **c**-**e** are HPLC-DAD experiments, where the *x*-axis gives the elution time in minutes and the *y*-axis is absorbance units at λ = 325 nm. The plots shown in **b** are UV spectra with the *y*-axis showing absorbance and the *x*-axis showing wavelength in nanometers.

Like other unmethylated, polyunsaturated pyrones, the enzymatic reaction products were challenging to isolate and characterize because of chemical instability. To gain further evidence for the presence of the pyrone structure, we hydrogenated and derivatized the enzymatic reaction mixture, and then analyzed the products by gas chromatography-mass spectrometry (GCMS). This method was chosen because of its predictive value: hydrogenation of the triene should produce four separable stereoisomers, with predictable fragmentation serving to localize methyl groups. Hydrogenation gave four isomers with nearly identical fragmentation patterns around some of the methyl groups, supporting the structural assignment (Extended Data Fig. 5).

Although some animal FAS/PKS enzymes accept methylmalonyl-CoA *in vitro*, malonyl-CoA is preferred^32–34^. We sought to determine whether EcPKS1 specifically incorporates methylmalonate, or whether it also uses malonyl-CoA like other enzymes. We incubated EcPKS1 with malonyl-, methylmalonyl-, acetyl-, and propionyl-CoA under various conditions (Fig. 3D, Extended Data Fig. 6). These experiments demonstrate that EcPKS1 uses only methylmalonyl-CoA, and not other substrates. Further, our data suggest that other CoA esters compete with methylmalonyl-CoA but are not incorporated into any products, indicating stringent selectivity for methylmalonate. *S*-adenosylmethionine (SAM) is the source of methyl groups in many branched polyketides^28^. We co-incubated SAM with malonyl-CoA and methylmalonyl-CoA in the presence of EcPKS1, showing that SAM is not involved in the biosynthesis (Extended Data Fig. 6). Thus, EcPKS1 is the first characterized methylmalonate-specific FAS/PKS from animals.

These data revealed that EcPKS1 is a mollusk polypropionate synthase. In combination with previous experimental evidence, a biosynthetic pathway from CO_2_ to complex polyketides can be defined. Symbiotic chloroplasts fix CO_2_, which is then converted into methylmalonyl-CoA. The animal enzyme EcPKS1 synthesizes the tetraene pyrone, which would be converted to tridachiones and related natural products by methylation, photorearrangement, and oxidation. These later events may be important in photosynthesis and are strongly supported by previous synthetic chemistry experiments in which the γ-methylated tetraene pyrone is used as starting material to synthesize the natural products (Extended Data Fig. 7)^35–37^.

The close connection between chloroplasts and polyene pyrones extends beyond biosynthesis. It has been widely observed that polyene pyrones are found in sacoglossans that engage in long-term photosynthesis, while those lacking polyene pyrones also derive little or no nutrition from photosynthesis (Extended Data Table 1). We found that, in *E. diomedea*, polyene pyrones are not found in developing embryos, which lack chloroplasts (Extended Data Fig. 8). Moreover, when examining transcriptomes of *E. chlorotica* at various growth stages, we found that *EcPKS1* is highly expressed as the animals metamorphose from the planktonic larval to the benthic juvenile slug stage, just before chloroplasts are acquired (Extended Data Fig. 9). The expression continues well into the stage when the animal maintains stable chloroplasts. In *E. timida*, expression of the *EcPKS1* ortholog *EtPKS1* is downregulated after prolonged starvation, when sequestered chloroplasts have degraded.

To examine the association of *EcPKS1* with photosynthetic ability further, we reinvestigated the *P*. cf. *ocellatus* and *E. diomedea* metagenomes with the hypothesis that *EcPKS1* homologs would be found in each, as these species are also capable of long-term chloroplast retention. Indeed, homologs of *EcPKS1, EcPKS2*, and cytoplasmic *FAS* were discovered (Extended Data Fig. 10). These were mollusk-encoded, as they came from intron-bearing contigs similar to those encoding their relatives in *E. chlorotica*. Our data thus strongly support previous suggestions that pyrones are critical for maintaining long-term photosynthetic activity in sacoglossans by serving antioxidant and photoprotective roles.

Polypropionates in marine mollusks are abundant and structurally diverse^13^. Aside from those described in this study, no other sequenced mollusks are known to contain polypropionates. Examining non-propionate mollusks in GenBank, we found that octopus, sea hares, clams, and oysters contain many FAS-type PKSs that have not been previously noted. However, none of these enzymes falls within the sacoglossan polypropionate group. So far two sequenced transcriptomes encode the methylmalonate-specific motif, from the hemichordate *Saccoglossus kowalevskii* and the beetle *Leptinotarsa decemlineata*. These data imply that the ability to synthesize complex polyketides is a ubiquitous feature among mollusks, and it may therefore play an important and unrecognized biological role in the animals.

Using these newly identified sequences, we generated a taxonomically broader phylogenetic tree based upon the ketosynthase domains of FAS/PKS genes (Extended Data Fig. 10). Many of the mollusk PKSs, as well as several from other phyla, fall in a single large group that includes animal FAS. A second large group contains all of the characterized type I PKS genes from bacteria, animals, and fungi. EcPKS1 is the only characterized PKS within the large FAS-like PKS cluster, representing a new biochemical frontier for PKS research.

A remaining mystery involves the *in vitro* product selectivity observed in our study. The major enzymatic product incorporated six units of methylmalonate, while the minor product incorporated seven. While many mollusk polypropionates contain five or seven equivalents of methylmalonate, none of the reported compounds contains six. We are therefore still missing a chain length-determining factor. EcPKS1 lacks a thioesterase (TE) domain, which is often important for chain length determination^38^. We attempted to use base to improve offloading of the product^39^, but were unsuccessful, indicating that an as-yet unknown factor governs chain length (Extended Data Fig. 6). We speculate that this factor is likely to be the unknown methyltransferase involved in the next step of the biosynthetic pathway.

In summary, EcPKS1 represents the first example of complex polyketide biosynthesis in animal metabolism. We use biochemical evidence to demonstrate that structurally elaborate polyketides, a family previously associated with microbial metabolism, are animal products as well. The biosynthesis is part of an elaborate molecular dance in which kleptoplasts from algae fix CO_2_. Fixed carbon is transformed into methylmalonyl-CoA and modified by the mollusk EcPKS1 enzyme to synthesize UV- and oxidation-blocking pyrones that protect the mollusk and its chloroplasts from photosynthetic damage.

## Methods

### Genomic DNA extraction, sequencing and assembly

Single live specimens of *Plakobranchus* cf. *ocellatus* “aff. sp. 1” sensu (collected by SCUBA in the Solomon Islands, June 22, 2006; 9° 1.02’S 160° 13.06’ E) and *Elysia diomedea* (purchased from www.Liveaquaria.com, Rhinelander, WI, USA) were used in this work. Tissue sections each approximately 0.5 cm^3^ were frozen using liquid nitrogen and ground using a sterile mortar and pestle. Ground tissue was resuspended in lysis buffer (2 mL; 100 mM NaCl, 10 mM EDTA, 1.0 % SDS, 50 mM Tris-HCl at pH 7.5) followed by addition of proteinase K (25 µL; 20 mg/mL) and incubated in a water bath at 55 °C until lysis was complete (clear solution). The solution was removed from the water bath and RNAse (2 µL; 20 μg/mL) was added. After 15 minutes of incubation at 37 °C, saturated KCl (200 µL) was added to the solution. The mixture was incubated in an ice bath for 10 minutes and then centrifugated at 3,739 x *g* at 4 °C. Phenol:chloroform extraction of the supernatant^40^ led to purified DNA. Sequencing was performed at the Huntsman Cancer Institute’s High Throughput Genomics Center, University of Utah. Sacoglossan genomes were sequenced using Illumina HiSeq 2000 sequencer with 350 bp inserts and 125 bp paired-end runs. Raw reads were merged using BBMerge^41^. Non-merged reads were filtered and trimmed using Sickle with the parameters (pe sanger -q 30 –l 40)^42^. The trimmed and merged FASTQ files were assembled using metaSPAdes^43^ with standard parameters in the Center for High Performance Computing at the University of Utah.

### Identification of polypropionate PKS genes in published sacoglossan transcriptomes

Sacoglossan transcriptome reads were acquired in previous work by other groups^11,23^ from 19 specimens of *Elysia chlorotica, Elysia timida*, and *Elysia cornigera* (Extended Data Table 3). We downloaded the reads from the NCBI Sequence Read Archive (SRA) and assembled them using IDBA-UD^44^. The assembled transcriptomes were queried by Basic Local Alignment Search Tool (BLAST) using the *Homo sapiens* FASN gene (NCBI Accession: NP_0045095.4) involved in human fatty acid metabolism. Putative full-length, assembled FASN-related transcripts were translated using ExPASy translation tool. FASN and related sacoglossan transcripts consist of multiple domains, which were annotated using Conserved Domain Database (NCBI)^45^ and Pfam 32.0 (EMBL)^46^. Geneious 10.2.2 software package was used to perform alignments and tree building. Sequence alignments were made using Multiple Sequence Comparison by Log Expectation^47^, trimmed and realigned. Phlyogenetic trees were constructed using the Unweighted Pair Group Method with the Jukes-Cantor model^48^.

### Construction of EcPKS1 expression plasmid

The plasmid was constructed by homologous recombination in yeast. The predicted *EcPKS1* gene was synthesized in three fragments containing 70-bp overlapping regions (Genewiz). The fragments were designed so that their ends overlapped with the promoter and terminator sites of yeast expression plasmid^39^. The expression vector was linearized using NdeI and PdeII (New England Biolabs) following the manufacturer’s protocol, and the *EcPKS1* fragments and linearized vector were cotransformed into *Saccharomyces cerevisiae* BJ5464 using the S.C. EasyComp Transformation Kit (Sigma). Transformants were plated onto synthetic complete media and selected by transformation to uracil prototrophy. Yeast colonies were combined, and plasmids were isolated using the QIAprep Spin Miniprep Kit (QIAGEN). Plasmids were amplified by transformation into chemically competent *Escherichia coli* DH5α (New England Biolabs), which was plated onto LB agar (10 g/L tryptone, 5 g/L yeast extract, 10 g/L NaCl, agar 20 g/L) with ampicillin (50 μg/mL) and incubated for 24 hours at 37 °C. Individual colonies were picked and inoculated in to 5 mL LB broth with ampicillin (50 μg/mL) and incubated at 30 °C with overnight shaking. Plasmids were obtained using the QIAPREP Spin Miniprep Kit (QIAGEN) and screened by digestion with BamHI (New England Biolabs) followed by agarose gel electrophoresis. Plasmid pJTPKS1 containing the expected insert was verified by Sanger sequencing.

### Overexpression and purification of EcPKS1

Plasmid pJTPKS1 was transformed into *Saccharomyces cerevisiae* BJ5464-NpgA (MATα ura3-52 trp1 leu2-Δ1 his3Δ200 pep::HIS3 prb1d1.6R can1 GAL), which was selected on uracil-deficient agar (1.39 g/L Yeast Synthetic Drop-out Media Supplements without uracil (Sigma-Aldrich), 6.7 g/L Yeast Nitrogen Base (Sigma-Aldrich), 40 mL/L 50% glucose solution, agar 20 g/L) and incubated at 30 °C for 48 hours. A single colony of BJ5464-NpgA-EcPKS1 was used to inoculate uracil-deficient broth (5 mL; 1.39 g/L Yeast Synthetic Drop-out Media Supplements without uracil (Sigma-Aldrich), 6.7 g/L Yeast Nitrogen Base (Sigma-Aldrich), 40 mL/L 50% glucose solution) and incubated at 30 °C with shaking at 150 rpm. After 24 hours, 1 mL of the culture was used to inoculate 6 × 1 L yeast peptone dextrose broth (10 g/L yeast extract, 20 g/L peptone, 20 g/L glucose). The culture was grown at 30 °C under constant shaking at 180 rpm. After 72 hours, the cells were pelleted at 3,739 × g for 20 minutes at 4 °C. The cell pellets were resuspended in lysis buffer (50 mM NaH_2_PO_4_, 150 mM NaCl, 10 mM imidazole, pH 8.0) and sonicated on ice at 1-minute intervals until homogenous. The resulting mixture was centrifugated at 28,928 × *g* for 40 minutes at 4 °C to separate the supernatant from the cell debris. The supernatant was filtered using a 0.45-micron PVDF syringe filter (Millex-HV, Sigma) before adding Ni-NTA and incubating the mixture at 4 °C for 12 hours. Soluble EcPKS1 in the lysis buffer was loaded onto Ni-NTA resin in a gravity column. The resin was washed with 25 mL each of 20 mM and 50 mM imidazole in 50 mM Tric-HCl buffer (500 mM NaCl, pH 8.0). The protein was eluted using 3 x 5 mL 250 mM imidazole in 50 mM Tris-HCl buffer pH 8.0. 15 mL of eluted EcPKS1 was concentrated to 200 μL, buffer exchanged (15 mL of 50 mM Tris-HCl, 2 mM EDTA, 5 mM DTT, pH 8.0) and further concentrated to 200 μL final volume using Amicon Ultra 100 MWCO centrifugal filters (EMD Millipore). Protein concentration was calculated to be 6.2 mg/L using the Bradford assay with bovine serum albumin (New England Biolabs) as a standard.

### *in vitro* characterization of EcPKS1

Optimization of the enzymatic reaction was done using 100-µL enzyme reaction mixtures consisting of 10 μM EcPKS1, 2 mM NADPH, 1 mM DTT and 2 mM methylmalonyl-CoA buffered using 100 mM sodium phosphate (over a range of pH 6.5-8.0) and incubated at room temperature under dark conditions. After 16 hours of incubation, reaction products were extracted twice with ethyl acetate (200 μL). The organic layer was dried using a speed vacuum concentrator and reconstituted with methanol and analyzed by HPLC using a Hitachi Primade HPLC system equipped with a PDA detector and autoinjector. Aliquots (20 μL) of the sample were injected onto a reversed-phase C18 analytical column (Luna C18 4.6 × 100mm, Phenomenex) with a gradient starting at 20% mobile phase B (H_2_O + 0.1 % TFA: MeCN) for 5 minutes and increased until 100 % mobile phase B for 20 minutes at a rate of 0.7 mL/min. The optimum working pH for EcPKS1 was determined to be 7.0 (Extended Data Fig. 6). HRMS analysis was performed at the University of Utah’s Mass Spectrometry and Proteomics Core using an Agilent 1290 equipped with an Agilent 1290 FlexCube and Agilent 6350 Accurate Mass Q-TOF dual ESI mass spectrometer. Sample (10 μL) was injected onto a UPHLC C18 column (2.1 × 150 mm Eclipse Poroshell, 1.9 μ ID) set at 30 °C. Elution was done using solvents containing 10 mM ammonium formate (AF) and 0.1% formic acid (FA). Mobile phase A contained water with AF and FA, while mobile phase B contained 95% acetonitrile (aqueous) with AF and FA. The gradient started at 10% mobile phase B for 1 minute, increasing to 100% B over 10 minutes. The flow rate was 0.5 mL/min. Electrospray ionization was done in positive mode using nitrogen as sheath gas at 350 °C flowing at 12 L/min. Capillary and nozzle voltages were set to 3500 V and 1000 V, respectively.

### EcPKS1 product offloading assay

Six 100-µL reaction mixtures containing 2 mM methylmalonyl-CoA, 1 mM DTT, 2 mM NADPH and 10 μM EcPKS1 in 100 mM phosphate buffered at pH 7.0 were incubated in the dark at room temperature for 16 hours. After incubation, 1 M NaOH (20 µL) was added to three of the reaction mixtures, which were then incubated at 65 °C for 10 minutes. Reaction products were extracted into ethyl acetate extraction analyzed by HPLC. For comparison, three control reaction mixtures were made using the same method, but not subjected to NaOH treatment (Extended Data Fig. 6).

### Methytransferase domain activity assay

Reaction mixtures (100-µL) containing 2 mM malonyl-CoA, 1 mM DTT, 2 mM NADPH, 5 mM S-adenosyl-L-methionine (SAM), 1mM MgCl_2_ and 10 μM EcPKS1 in 100 mM phosphate buffer (pH 7.0) were incubated in the dark at room temperature. Two parallel enzymatic reaction mixtures containing additional 2 mM methlymalonly-CoA and one without SAM and MgCl_2_ but with 2 mM methylmalonyl-CoA were carried out for comparison. After 16 hours of incubation, reaction products were extracted with two volumes of ethyl acetate method and analyzed using HPLC (Extended Data Fig. 6).

### Acyl substrate specificity assays

Acyl-CoA substrates malonyl-Co A, acetyl-CoA, propionyl-CoA, and methylmalonyl-CoA were purchased from CoALA Biosciences. Enzyme reaction mixtures (100 μL) consisted of enzyme EcPKS1 (10 μM), sodium phosphate pH 7.0 (100 mM), NADPH (2 mM), DTT (1 mM) and acyl-CoA substrates (2 mM). When multiple acyl-CoA substrates were used in a single reaction, they were used at 2 mM each. Reaction mixtures were incubated at room temperature for 16 hours and protected from light to avoid photochemical rearrangements (Extended Data Fig. 6).

### Chemical characterization of enzyme reaction products

To generate EcPKS1 products for chemical characterization, fifty enzyme reaction mixtures (100-µL) containing 2 mM methylmalonyl-CoA, 1 mM DTT, 2 mM NADPH and 10 µM EcPKS1 in 100 mM phosphate buffer (pH 7.0) were incubated at room temperature for 48 hours with addition of 200 mM NADPH (2 µL) to replenish NADPH after the 24^th^ hour. Enzyme reaction mixtures were pooled and loaded into C18 resin packed in cotton-plugged borosilicate glass Pasteur pipets. The resin was washed twice with water (1 mL) followed by 20% acetonitrile:water (1 mL) before elution with acetonitrile (3 mL). The eluant was dried using a speed vacuum concentrator and reconstituted in methanol 400 µL. Pd/C (0.5 mg) was added to the solution and stirred for 16 hours under a H_2_ atmosphere. The mixture was then filtered through Celite to remove Pd/C and dried a using speed vacuum concentrator. GC-MS/MS analysis was performed at the University of Utah’s Metabolomics Core.

GC-MS analysis used an Agilent 7200 GC-QTOF and an Agilent 7693A automatic liquid sampler. Dried samples were suspended in dry pyridine (100 µL; EMD Millipore) containing *O*-methoxylamine hydrochloride (40 mg/mL; MP Bio) and incubated for one hour at 37 °C in a sand bath. An aliquot (50 µL) of the solution was added to autosampler vials. *N*-Methyl-*N*-trimethylsilyltrifluoracetamide containing 1% chlorotrimethylsilane (60 µL; Thermo) was added automatically via the autosampler and incubated for 30 minutes at 37 °C. After incubation, samples were vortexed and the prepared sample (1 µL) was injected into the gas chromatograph inlet in the split mode with the inlet temperature held at 250 °C. A 1:1 split ratio was used for analysis. The gas chromatograph had an initial temperature of 60°C for one minute followed by a 10 °C/min ramp to 325 °C and a hold time of 10 minutes. A 30-meter Agilent Zorbax DB-5MS with 10 m Duraguard capillary column was employed for chromatographic separation. Helium was used as the carrier gas at a rate of 1 mL/min.

Polyene (**1**): UV (MeCN) λ_max_ 279, 318 nm. HRESIMS *m/z* 289.1814 [M+H] ^+^ (calcd for C_18_H_24_O_3,_ 289.1798). GCMS/MS of **1c**: Parent mass *m/z* 366.2590 (calcd for C_21_H_38_O_3_Si, 366.2548), Fragments: *m/z* 73.0475 (calcd for C_3_H_9_OSi, 73.0474), *m/z* 155.0873 (calcd for C_11_H_23_, 155.1800), *m/z* 211.1141 (calcd for C_20_H_35_O_3_Si, 211.0790), *m/z* 253.1239 (calcd for C_13_H_21_O_3_Si, 253.1260), *m/z* 351.2322 (calcd for C_20_H_35_O_3_Si, 351.2355).

Tridachione precursor (**2**): UV (MeCN) λ_max_ 260, 355 nm. HRESIMS *m/z* 329.2120 [M+H] ^+^ (calcd for C_21_H_28_O_3_, 329.2111).

### Chemical analysis of *E. diomedea*

Chemical extracts were prepared by homogenizing egg ribbons and tissue sections (approximately 10 cm^3^) of *E. diomedea* with acetone (5 mL). The homogenate was centrifuged at 3,739 x *g* for 20 minutes at 4 °C and the supernatant recovered by aspiration. The acetone extract was dried using a rotovap and resuspended in methanol. Extracts were analyzed using a HPLC using a Hitachi Primade HPLC system equipped with a PDA detector and autoinjector. Aliquots (20 µL) were loaded onto a C18 analytical column (Luna C18 4.6 x 100mm, Phenomenex) with a gradient starting at 30% mobile phase B (H_2_O + 0.1 % TFA: MeCN) and 70% mobile phase A (H_2_O) for 5 minutes and increased until 100 % mobile phase B for 25 minutes at a rate of 1.0 mL/min. Purification of the major polypropionate in *E. diomedea* tissue extract was performed using a semi-preparative HPLC column (Luna 5u C18 250 x 10 mm) at a flowrate of 2.5 mL/min. ^1^H NMR data were obtained using a Varian 500 (^1^H 500 MHz) NMR spectrometer equipped with a 3mm Nalorac MDBG probe, utilizing residual CDCl_3_ signal for referencing. Natural products, including tridachione, were identified by comparing data to those reported in the literature. A similar procedure was used to isolate and identify products from *P*. cf. *ocellatus*.

### Discovery of EcFAS, EcPKS1, and EcPKS2 analogs in *P*. cf. *ocellatus* and *E. diomedea*

Gene prediction was done using AUGUSTUS^49^. Full-length *EcPKS1, EcPKS2*, and *EcFAS* genes were used as queries to search for PKS gene-containing contigs by tblastn search (E-value = 1.0 × e^-10^) in *P*. cf. *ocellatus* and *E. diomedea* genomic assembly. Contig hits were then submitted to AUGUSTUS to predict genes based on parameters built using the reported *E. chlorotica* genome assembly and protein sequences^50^. The resulting predicted gene fragments were queried against EcPKS1, EcPKS2 and EcFAS using blastp search (E-value = 1.0 × e^-10^) and were ranked according to bitscore and E-value to determine the best hit. The best hits for EcPKS1, EcPKS2 and EcFAS were assigned to each query gene then aligned and assembled to create the predicted PKS and FAS orthologs in *P*. cf. *ocellatus* aff. sp. 1 and *E. diomedea* (PoPKS1, PoPKS2, PoFAS for *P*. cf. *ocellatus* aff. sp. 1 and EdPKS1, EdPKS2 and EdFAS for *E. diomedea*).

### EcPKS1 gene expression analysis

Raw reads were trimmed by Sickle with parameters (pe sanger –q 30 –l 40). Trimmed reads were pooled by species, and *de novo* assembly for each species was done (EC and ET) using maSPAdes with default parameters. Trimmed raw reads of each specimen were multimapped to the reference transcriptome assembly using Salmon^51^ with parameters (salmon index –t –i index –k 31; salmon quant --index –valudateMappings –libType A –dumpEq –r). The multitude of contigs produced by assembly were hierarchically clustered, and cluster count summarization was performed using Corset 1.04^52^ according to shared read information with parameters (-f true –g –n – salmon+eg_classes). Transcript abundance of clusters (EcPKS1 gene) was estimated using EdgeR^53^ by normalized counts per million.

## Data and materials availability

Plasmids are available from the corresponding author with a standard MTA. All sequences described in this study have been deposited in GenBank (See Extended Data Table 3 for accession numbers).

## Acknowledgements

We thank James Cox and John Alan Maschek from the University of Utah Metabolomics Core Facility, which is supported by 1 S10 OD016232-01, 1 S10 OD021505-01 and 1 U54 DK110858-01. Yi Tang kindly provided *S. cerevisiae* BJ5464-NpgA. Paul Jones from Wake Forest University and John Moses from LaTrobe University provided helpful discussion and advice concerning the chemistry of mollusk pyrones. We are grateful to the people and nation of Solomon Islands for permission to perform the study on *P. ocellatus* following Nagoya Protocol standards. J.P.T., Z.L., and E.W.S. are supported by NIH R35GM122521 and U01TW008163.

## Contributions

J.P.T. and E.W.S. conceived of the project. J.P.T. performed experiments and bioinformatics to discover EcPKS1. Z.L. performed bioinformatics. J.M.W. provided assistance and guidance concerning yeast expression and biochemical characterization experiments. P.J.K. helped to interpret the results and helped with mollusk taxonomy. E.W.S. and J.P.T. wrote the manuscript with input from all authors.

## Corresponding author

Correspondence to Eric W. Schmidt.

